# Spectral graph model for fMRI: a biophysical, connectivity-based generative model for the analysis of frequency-resolved resting state fMRI

**DOI:** 10.1101/2024.03.22.586305

**Authors:** Ashish Raj, Benjamin S Sipes, Parul Verma, Daniel H. Mathalon, Bharat Biswal, Srikantan Nagarajan

## Abstract

Resting state functional MRI (rs-fMRI) is a popular and widely used technique to explore the brain’s functional organization and to examine if it is altered in neurological or mental disorders. The most common approach for its analysis targets the measurement of the synchronized fluctuations between brain regions, characterized as functional connectivity (FC), typically relying on pairwise correlations in activity across different brain regions. While hugely successful in exploring state- and disease-dependent network alterations, these statistical graph theory tools suffer from two key limitations. First, they discard useful information about the rich frequency content of the fMRI signal. The rich spectral information now achievable from advances in fast multiband acquisitions is consequently being under-utilized. Second, the analyzed FCs are phenomenological without a direct neurobiological underpinning in the underlying structures and processes in the brain. There does not currently exist a complete generative model framework for whole brain resting fMRI that is informed by its underlying biological basis in the structural connectome. Here we propose that a different approach can solve both challenges at once: the use of an appropriately realistic yet parsimonious biophysical signal generation model followed by graph spectral (i.e. eigen) decomposition. We call this model a Spectral Graph Model (SGM) for fMRI, using which we can not only quantify the structure-function relationship in individual subjects, but also condense the variable and individual-specific repertoire of fMRI signal’s spectral and spatial features into a small number of biophysically-interpretable parameters. We expect this model-based inference of rs-fMRI that seamlessly integrates with structure can be used to examine state and trait characteristics of structure-function relations in a variety of brain disorders.

## 1 Introduction

Understanding the governing principles underlying the brain’s resting state activity is an area of immense importance. Since it’s first documentation in 1990 [1], the blood oxygenation level dependent (BOLD) signal in functional magnetic resonance imaging (fMRI) has been a critical and quickly evolving method to study changes in brain activity during task and rest. More recently, the synchronized fluctuations between brain regions have been characterized as a type of functional connectivity (FC) [2]. This has itself launched yet another paradigm shift toward analyzing BOLD signals as a network, where connections across regions are defined as the Pearson’s correlation between the average BOLD signal time series of those respective regions [3]. This network science framework has opened lines of inquiry that seek to model and therefore predict empirical FC using only the brain’s wiring diagram, known as it’s structural connectivity (SC), as measured by diffusion-weighted MRI. Despite these successes, resting state (rs-)fMRI analysis methods have reached saturation, and the next generation of techniques will need to address two broad limitations: the focus on the fMRI signal’s covariance (FC) without considering its power spectrum; and the use of statistical rather than biophysical models to obtain features of interest. Since every limitation provides an opportunity for a paradigm shift, below we describe the opportunities available for future methodological innovation. We then introduce our proposal, which we believe could be an early but important step in this direction.

### Opportunity to accommodate both FC and spectral content

Current methods broadly seek to model, analyze, or extract features of interest from functional connectivity (FC) - i.e. the 2nd order covariance structure inherent in the data. Whether this FC is evaluated via pairwise Pearson’s correlation between time series, or through ICA to achieve functional connectivity networks (FCNs), it essentially neglects the actual temporal structure of fMRI time series, specifically their frequency content. The BOLD signal is generally bandpass-filtered from 0.01–0.08 Hz to avoid contamination by physiological frequency artifacts; this practice is so common that this frequency filter range is set by default in popular connectivity processing pipelines [4, 5]. Historically, this has not been a particular encumbrance due to the slow nature of BOLD response and the long TRs needed for full brain coverage. Yet consequentially, the spectral content of the fMRI BOLD signal has remained under-addressed.

There is mounting evidence, however, that higher frequencies in spontaneous BOLD fMRI activity may be meaningful. Using advanced accelerated fMRI techniques, canonical resting-state networks (RSNs) have been reported with components at higher frequencies up to 1.5 Hz [6–8]. Further work has shown that BOLD frequency bands throughout the range 0.01–0.25 Hz were highly reproducible and had meaningfully varying network topology across RSNs [9]. While caution is still needed regarding high-frequency BOLD noise contamination [10], de-noising techniques that remove physiological noise sources demonstrate impressive and reproducible results [11, 12], thus opening the door to meaningful high-frequency BOLD analysis. It is already evident that certain frequency-specific measures, like the fraction of low-frequency power, are useful descriptors of diseases like Schizophrenia [13]. Phase-sensitive coherence was previously proposed as a measure of FC, convincingly demonstrating the value of frequency-resolved FC [14, 15].

Although generative models of fMRI are available via dynamic causal modeling (DCM), which infer the coefficient matrix of an underlying vector autoregression (VAR) process [16], they were historically limited to correlation-based FC and not power spectra. Recent advances in spectral dynamic causal modeling has expanded their ability to parameterize fMRI coherence spectra with shape and scale parameters [17, 18]. This is an important advance but one that remains phenomenological and unrelated to the underlying structural substrate or biophysical processes.

### Opportunity to move from statistical to biophysical descriptors of fMRI

Second, current methods are to a large extent statistical - they seek to uncover the difference in summary graph theoretic statistics of FC matrices between diagnostic or task groups. A large body of work has evolved to uncover the graph theoretic features of brain FC networks. These sophisticated network analysis methods have proven value in differentiating between disease [19, 20] and cognitive domains [21], and have proved especially helpful in exploring the neural correlates of developmental and psychiatric disorders [22]. Such methods are indeed highly valuable from a practical point of view, but do not typically produce new insight about the underlying biological or biophysical processes that give rise to the observed fMRI data. Despite their broad success, network methods by design remain limited to statistical rather than biological descriptors of brain activity. Indeed, the broad success of graph analysis methods suggests that such a biophysical interpretability is not necessary for many application areas. Yet, the ability to connect the underlying biophysics with observed FC is not only possible but may also lead to enhanced understanding of brain phenomena.

The availability of frequency-rich fMRI further enhances the opportunity to achieve physiologically-relevant analysis, as higher frequencies may arise from a combination of the hemodynamic response, neural and cortical time constants, and the aggregate behavior of the brain’s slow oscillatory dynamics and global coupling. Hence there is an opportunity to explore biophysically-grounded models of how fMRI time series, its spectral content, and especially its higher frequencies, arises from the brain’s structural substrate. Previous biophysical models of neural activity on coupled systems [23, 24] have been very successful in capturing empirical fMRI signal’s FC but were not designed to capture its spectral content – see Discussion.

### Proposing a biophysical model-based approach to analyze wideband rs-fMRI spectra and frequency-dependent FC

Arguably, both opportunities identified above may be addressable via biophysically realistic yet appropriately parsimonious signal generation models that can interrogate BOLD frequency content and its frequency-dependent FC simultaneously. In this article, we propose a different approach of analyzing fMRI data via a powerful yet simple, parsimonious, and linear signal generation model arising from the underlying structural substrate. We show that a full mathematical exposition of this model-based approach can parsimoniously and elegantly exploit both opportunities listed above. Remarkably, this generative model is solved directly in frequency domain *in closed form* as a summation over graph eigenspectra. Hence we call this model a Spectral Graph Model (SGM) for fMRI. It can capture both wideband spectral as well as second-order covariance structures (i.e. frequency-resolved cross-spectral density) simultaneously. We implemented statistical inference to obtain optimal parameters fitted to two large cohorts of healthy subjects. Our approach obviates the need for large time-consuming simulations or complicated inference procedures.

The fitted model thus constitutes not only a mapping between structure and function in the brain, but also gives a novel way of analyzing spectrally-rich fMRI data. Furthermore, the biologically meaningful global parameters may offer crucial and individualized context to how the subject’s fMRI signals, their spectra and network topology are related to the underlying structural wiring substrate. We expect this model-based analysis tool will aid practitioners in analyzing fMRI data in a novel, frequency-resolved environment. An analysis overview is depicted in Figure 1.

**Figure 1:**
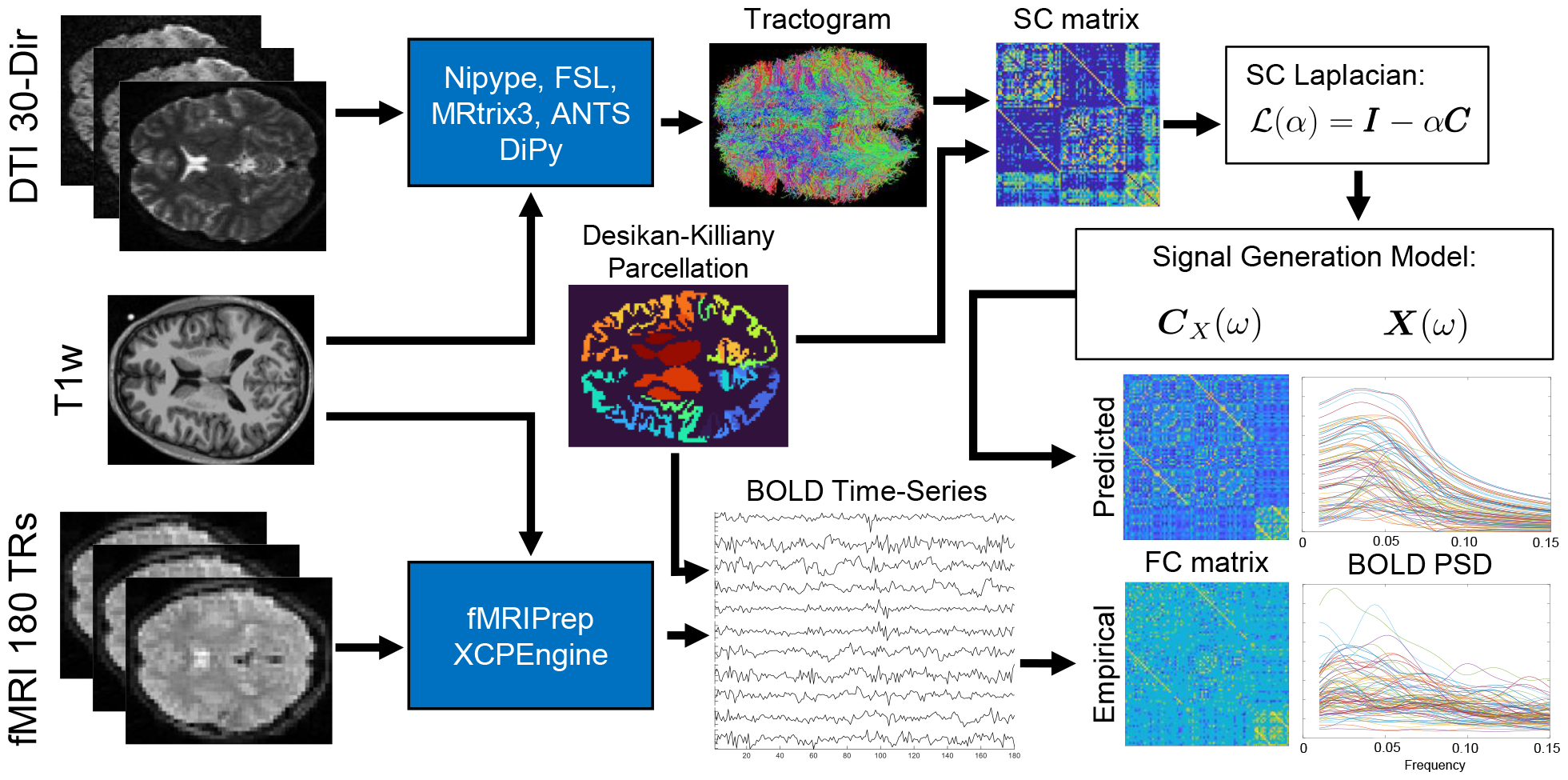
Analysis Overview. We implemented two processing streams, one for structural connectivity (SC) and one for functional connectivity (FC). The 30-direction diffusion weighted MRI images along with T1w structural images were preprocessed within a Nipype pipeline using FSL and MRtrix3. GPU-accelerated DiPy generated whole-brain seeded probabilistic tractograms, which were input to a post-processing stream with MRtrix3 and ANTS. The SC matrix (*C*) was defined as the probabilistically weighted number of streamlines between regions defined by the Desikan-Killiany (DK) atlas. Resting-state T2*w images with 180 timepoints were preprocessed with fMRIPrep and post-processed with XCPEngine to generate region-wise BOLD time series, also partitioned by the DK atlas. We sought to predict two aspects of the functional time series: its FC matrix, defined as the Perason’s correlation between regional time series, and the power spectral density (PSD) of each region’s time series. SC predicts both FC PSD through a signal generation model that uses the eigenvalues and eigenvectors of the SC’s Laplacian (ℒ) and two global parameters, *α* and *τ*, which are optimized for each subject’s empirical FC and PSD simultaneously.

## 2 Mathematical Model

**Notation**. In our notation, vectors and matrices are represented in **bold**, and scalars by normal font. We define a vector of ones as **1**. The Fourier transform of a signal *x*(*t*) is denoted as *ℱ*{*X*}(*ω*) for angular frequency *ω* = 2*πf*, where *f* is the frequency in Hertz. The structural connectivity matrix is denoted by ***C*** = {*c*_*l,m*_}, consisting of connection strength *c*_*l,m*_ between any two brain regions *l* and *m*.

### 2.1 Generative model of Network Spread of fMRI signal

For an undirected, weighted graph representation of the structural network *C* = {*c*_*l,m*_}, we model the average BOLD fMRI activation signal for the *l*-th region as *x*_*l*_(*t*):

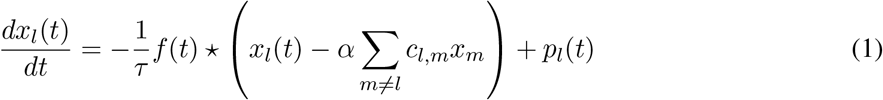

whereby the fMRI signal at the *l*-th region is controlled by a characteristic time constant *τ*, and input signals from regions *m* connected to region *l* are scaled by the connection strengths from *c*_*l,m*_. The symbol ⋆ stands for convolution, *p*_*l*_(*t*) is input noise driving the local signal at node *l*, and parameters *τ* is the characteristic time constant (in seconds), and *α* is a global coupling constant (unitless). These are global parameters and are the same for every region *l*. Below we describe each component of this model.

### 2.2 Impulse response *f*(*t*) and its time constant *τ*

The function *f* (*t*) represents the ensemble average neural impulse response, which accounts for various delays introduced by neural processes including synaptic capacitance, dendritic arbors, axonal conductance, and other local oscillatory processes that involve detailed interactions between excitatory and inhibitory neurons, etc. This impulse response is modeled by a Gamma-shape function with a single characteristic time constant, given by *τ* :

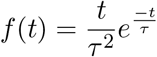

This impulse response may also be considered to incorporate the neurovascular coupling represented conventionally by the hemodynamic response function. Note that while previous analogous rate models for MEG frequencies explicitly introduce axonal conduction delays between brain regions [25, 26], here we instead incorporate those delays implicitly within *f*(*t*) via its single parameter *τ* rather than explicitly, since our specific focus is to explore much longer time lags in functional activity that arise from other sources. In the fMRI regime these various processes are not possible to separate or identify, hence a combined model parameter via *τ* is reasonable.

For convenience and parsimony we use the same time constant *τ* to also capture the self-decay term in the signal model (first term on the right hand side). This choice is deliberate, since the self-decay is intimately related to the local impulse response. We note that it would be straight-forward to allow two different time constants, one for the self-decay term and another for the impulse response. The effect of this choice was empirically explored in Results section.

### 2.3 Global coupling constant *α*

The global coupling parameter *α* acts as a controller of weights given to long-range white-matter connections. This parameter represents aspects of global integration and segregation, and may be mediated by attention, neuromodulatory systems and corticothalamic control signals. Again, these processes are not possible to be separately identified in the fMRI regime, hence lumping them makes sense. The above model then is the signal generation equation for fMRI.

### 2.4 Vectorizing via Laplacian Matrix

Let us define the Laplacian matrix

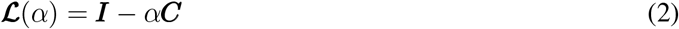

Where ***I*** is the identity matrix and ***C*** is the connectivity matrix as defined above. Then the above pair-wise equation readily generalized to the entire brain network, yielding a vectorized differential equation

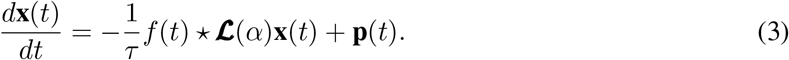

### 2.5 Reduction to the eigen-mapping model

We note first that 3 is a generalization of the passive diffusion model of brain activity spread–an idea that has been successfully applied to predict steady-state zero-lag FC from the SC’s Laplacian [27]. Indeed, removing the impulse response *f*(*t*), the above system admits a closed-form solution of the free-state evolution of the expected signal with initial condition **x**_0_ and no external driving signal **p**:

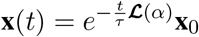

Following the framework well-described in [27, 28], the above signal equation readily yields the theoretical functional connectome from the Laplacian **ℒ** of the structural connectome as follows. Assume that only a single region *i* is experiencing activity at *t* = 0, hence **x**_0_ = **e**_*i*_, a unit vector with zeros everywhere except at the *i*-th location. Concatenating this for all regional activations in turn, we obtain:

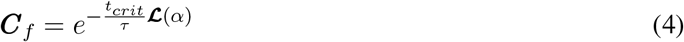

for some constant *t*_*crit*_ that may be empirically inferred. In our prior work [29, 30], this model was generalized to the “eigen-decomposition” model that relates the SC and FC matrices’ eigenvalues and eigenvectors; and further generalized to a series expansion of eigenvalues [31–35].

### 2.6 Novel Fourier-domain model of fMRI signal

Here we wish to introduce a richer signal model with local impulse responses, and impart the model with the ability to manifest rich frequency dependencies. Hence we propose the following linear systems approach that relies on Fourier transform of the signal equation. Since the above equations are linear, we can obtain a closed-form solution in the Fourier domain as demonstrated below.

Due to its Gamma shape the ensemble average neural response function *f*(*t*) admits a simple closed-form Fourier transform: 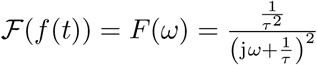 Taking Fourier transform of the vectorized signal equation (3), using the notation ℱ(**x**(*t*)) = **X**(*ω*) and ℱ(**p**(*t*)) = **P**(*ω*), we obtain:

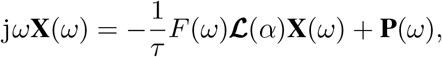

Here we used the Fourier pair 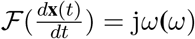. The above equation can be re-arranged as:

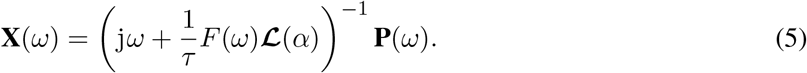

To evaluate this expression we employ the eigen-decomposition of the Laplacian:

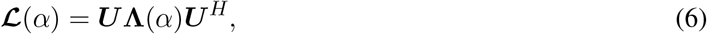

where, ***U*** = {**u**_*k*_, ∀*k* ∈ [1, *N*]} are the eigenvectors and **Λ**(*α*) = diag([*λ*_1_(*α*), …, *λ*_*N*_ (*α*)]) contains on its diagonal the eigenvalues for a given coupling constant *α*. We note that due to the definition of the Laplacian above, the eigenvectors do not have any *α* dependence, while the eigenvalues do.

Then the above signal equation can be re-written in closed form as the summation over the eigenvectors:

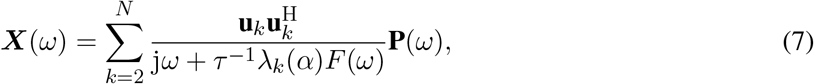

Equation (7) is the closed-form steady state solution of the macroscopic signals at a specific angular frequency *ω*. Henceforth we drop the explicit dependency on *α* whenever convenient. When this equation is required to be explicitly evaluated, we assume that ***P***(*ω*) = **1**, a vector of ones. This corresponds to either uniform stimulation or perturbation with white noise.

### 2.7 Novel frequency-resolved model of FC

With a signal generation equation in both time- and frequency domains, it is now possible to explicitly write the structure-function relationship in terms of the eigendecomposition of the structural Laplacian ℒ There are several equivalent ways to achieve this; here we use the most intuitive approach. Since the cross-spectral density (CSD) is given by ***C***_*X*_(*ω*) = ℰ(***X***(*ω*)***X***^*H*^(*ω*)), we get, under the assumption that ℰ(***P*** (*ω*)***P*** ^*H*^(*ω*)) = *σ*^2^***I***:

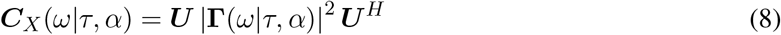

In this formulation the eigenvectors of the predicted FC are the same as those of the structural Laplacian (i.e. ***U***), while the eigenvalues are related via the diagonal matrix of new (frequency-dependent) eigenvalues **Γ**_*k,k*_(*ω*|*τ, α*) = *γ*_*k*_(*ω*|*τ, α*). In this manner we have reduced the full cross-spectral density of fMRI to modeling just the diagonal eigenvalues of the structural connectome; all region-pair coherences are thus expected to be captured entirely by the eigenvectors ***U***. The explicit relationship between the theoretical FC’s eigenvalues and those of the structural Laplacian’s is given by

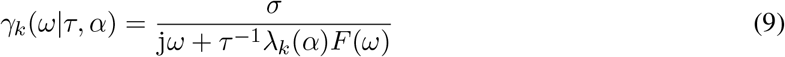

In this paper we will fit for both the theoretical signal spectrum (Eq 7) over all frequencies; and theoretical FC derived from the cross-spectral density (Eq 8).

## 3 Methods

### 3.1 Data acquisition and processing (UCSF cohort)

Data were collected as part of a multi-site longitudinal study aimed at better understanding the brain mechanisms underlying psychosis development and provided by our collaborators in the Brain Imaging and EEG Laboratory at the San Francisco VA Medical Center. Scanning was completed at the UCSF Neuroimaging Center using a Siemens 3T TIM TRIO with the following parameters

#### High-resolution structural T1-weighted images

MPRAGE, repetition time (TR) = 2,300 ms, echo time (TE) = 2.95 ms, flip angle = 9 degrees, field of view (FOV) = 256 x 256, and slice thickness = 1.20 mm

#### Resting fMRI

T2*-weighted AC-PC aligned echo planar imaging (EPI) sequence: TR = 2,000 ms, TE = 29 ms, flip angle = 75, FOV = 240 x 240, slice thickness = 3.5 mm, acquisition time = 6:22.

#### Diffusion-weighted MRI

b = 800s/mm2, 30 diffusion sampling directions, TR = 9,000 ms, TE = 84 ms, FOV = 256 x 256, and slice thickness = 2.00 mm. For the purpose of this study, we included only healthy subjects from the UCSF study who had both fMRI and DWI data, yielding 56 subjects (20 women; 23.8 *±* 8.3 years), and analysed only their baseline scans.

#### 3.1.1 Anatomical and Functional Preprocessing

The T1-weighted (T1w) images and T2*-weighted BOLD images were preprocessed using default procedures in fMRIPrep [36], which is based on Nipype [37] and Nilearn [38]. For more details on this pipeline, see the section corresponding to workflows in *fMRIPrep*’s documentation. For completeness, we summarize the anatomical and functional preprocessing below.

T1-weighted (T1w) images were corrected for non-uniformity [39, 40] and were used as a reference image throughout the workflow. The T1w-reference was skull-stripped and segmented based on cerebrospinal fluid, white-matter, and gray-matter [40, 41]. The T1w reference and template were spatially normalized to a standard space (MNI152NLin2009cAsym) with non-linear registration [40].

FMRIPrep generated a BOLD reference image that was co-registered to the T1w reference using flirt and boundary-based registration [42, 43]. Head-motion parameters were estimated before spatio-temporal filtering using FSL’s `mcflirt` [44]. BOLD runs were slice-time corrected and re-sampled onto their original, native space by applying transforms to correct for head motion [45]. The BOLD time-series were re-sampled into the same standard space as the T1w image (MNI152NLin2009cAsym). ICA-AROMA performed automatic detection of signal noise components, including motion artifacts, and saved for use during functional network generation [46].

#### 3.1.2 Functional network generation

Average functional time series were extracted from 86 regions of interest (68 cortical, 18 subcortical) as defined by the Desikan-Killiany atlas [47]. Functional connectivity processing followed the ICA-AROMA with global signal regression (GSR) pipeline as described in a benchmarking paper [48]. This pipeline included smoothing with a 6mm FWHM SUSAN kernel and regional time series bandpass filtering with a Butterworth filter from 0.01 Hz to 0.25 Hz.

Entries of (zero-lag) functional connectivity matrices were defined as the Pearson’s correlation between regional time series. We also obtained the full cross-spectral density (CSD) of the time series, denoted by the frequency-resolved matrix ***C***_*S*_(*ω*). Functional connectivity matrices were thresholded at the percolation threshold, which reduces the influence of noise in the network and maximizes the networks’ information relative to null models that preserve the strength distribution, degree distribution, and total weight [49, 50].

Because the percolation threshold seeks to maximize the network’s information in each FC, every individual had a subject-specific percolation threshold applied to their FC, which was used for model fitting.

#### 3.1.3 Structural network generation

Raw diffusion MRI data were processed with T1-weighted (T1w) images using Nipype [51], which implemented functions from FSL, MRtrix3, and ANTS. The raw DWI and T1w images were reoriented and registered to MNI space. T1w image brain extraction was performed with FSL BET [52]. DWI were denoised with MRtrix3 ringing removal, DWI bias correction, and Rician noise correction [53]. Fractional anisotropy in each DWI voxel was estimated from a fit tensor. Streamlines were probabilistically generated with whole-brain seeding using DiPy and an NVIDIA GPU [54]. Spherical-deconvolution informed filtering of tractograms (SIFT-2) determined the cross-sectional area multiplier for each streamline such that the streamline densities in each voxel are close to the fiber density estimated using spherical deconvolution [55]. The Desikan-Killiany atlas with 86 regions was linearly and non-linearly registered to DWI space using ANTS with GenericLabel interpolation, which parcellated streamlines into region-to-region structural connectivity [40]. Structural connectivity between regions was quantified as the probabilistic and SIFT-2 weighted number of streamlines between regions, normalized by the sum of all matrix entries. Additionally, we added inter-hemispheric and regional structural adjacency information to the structural connectome, then re-normalized by the sum of all entries, according to [56].

### 3.2 Public fast fMRI dataset (MICA cohort)

We sought to evaluate the model’s robustness and capacity to analyze fMRI data with richer frequency content. To this end, we used the recently published publicly available dataset for Microstructure-Informed Connectomics (MICA-MICs) with 50 healthy human subjects (23 women; 29.54 *±* 5.62 years) [57]. These data use similar pipelines to those described above (see [58] for processing details), but importantly, the fMRI data was collected with a 600 millisecond TR, meaning that the time series contains much richer frequency content. Since these data are without global signal regression (GSR), we performed GSR by extracting and removing the signal’s first principal component from all regions.

### 3.3 Practical Considerations Around Parameter Inference for Individual Subjects

While the theoretical model is simple and straightforward to evaluate simultaneously on FC and spectra, several implementation details were found to improve the quality and reliability of the fits to individual subjects. As described below, not all eigenmodes in the theoretical model are equally important, and not all portions of the signal cross-spectral density are equally useful.

#### 3.3.1 Alternate definitions of Spectral power and FC

We wish to achieve match between the theoretical cross-spectral density matrix ***C***_*X*_(*ω*|*τ, α*) in Eq 8, and the corresponding empirical CSD denoted by ***C***_*S*_(*ω*). However, a direct fitting to the 3D CSD array (*regions* × *regions* × *ω*) was found to be problematic: as depicted in Figure 2, both the empirical and theoretical CSD are extremely sparse matrices, with near-zero values at frequencies higher than around 0.10 Hz and especially in off-diagonal entries, but with a significant amount of noise coming from both the measured signal as well as highly challenging Fourier transform-related spectral estimation issues. Thus fitting to the entire CSD volume is extremely poorly posed. Therefore we developed an alternate strategy whereby we identified the most signal-rich and reliable portions of the CSD volume - the zero-lag FC (given by the integration over all frequencies of CSD: ∫ ***C***_*X*_(*ω*)*dω*); or a single slice of CSD ***C***_*X*_(*ω*_0_) at a fixed frequency *ω*_0_, the frequency where the fMRI spectral response was assessed to be the maximum for a given subject. This approach allows the model fits to be informed by the most critical elements of interest to the modeler, without necessarily suffering from the challenges explained above.

**Figure 2:**
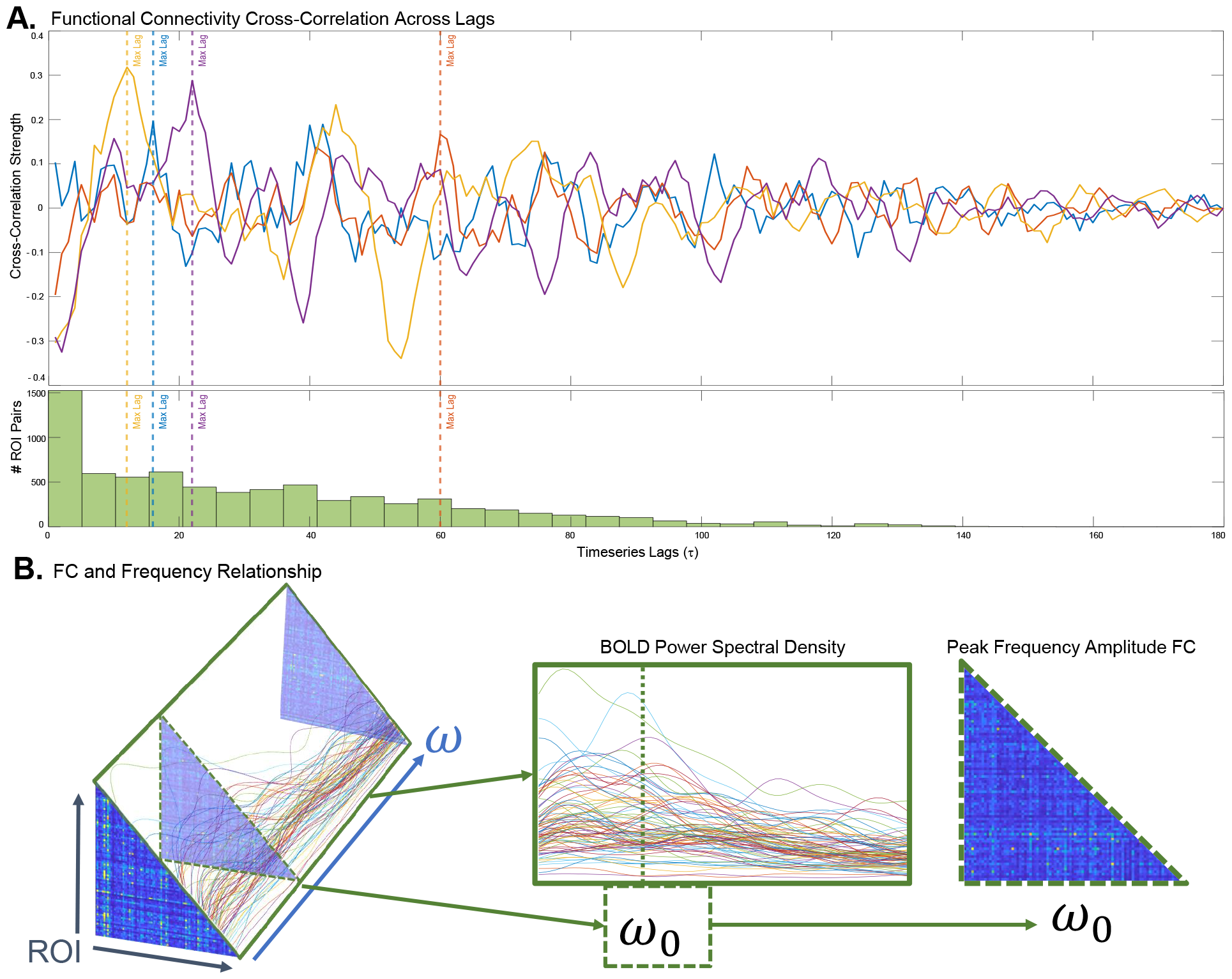
Frequency-resolved fMRI features contain meaningful information beyond the conventional zero-lag FC. The curves in panel **A** represent the cross-correlation between any two regions, revealing that the maximum cross-correlation frequently occurs at non-zero lags, and the accompanying histogram suggests a wide distribution of lags at which maximum correlation occurs. Since the Fourier Transform of the cross-correlation matrix is the cross-spectral density (CSD), the latter exhibits a frequency-selective distribution (panel **B**, left). In this study we carefully chose the most information-rich features from the full CSD: the power spectral density (PSD, defined as the diagonal of the CSD volume), middle panel; and CSD at a specific frequency *ω*_0_ at which the pairwise sum of CSD has the highest value for a given subject - right panel. The proposed structural connectivity-based SGM analysis aims to reproduce all these features simultaneously.

In similar vein, apart from the formal definition of PSD as the diagonal of the CSD, i.e. *diag*(***C***_*X*_(*ω*)), it is also useful to define simply the power of the signal under an explicit input, i.e. as |***X***(*ω*)|, when ***P*** (*ω*) = **1**.

#### 3.3.2 Eigenmode Weighting Algorithm

Not all eigenmodes in the summation shown in Eqn 7 are equally important or relevant for predicting FC. As others have noted, the SC matrix yields many eigenmodes that either do not participate in dynamic FC [59, 60], or do not contribute to models fitted to fMRI [56, 29] or MEG data [61]. Hence we developed a simple technique for evaluating eigenmode weighting by projecting the FC into the Laplacian’s eigenspace, thus performing a graph Fourier transform of the FC matrix: ***Q*** = ***U*** ^*T*^ ***F C U***. The diagonal entries of this matrix correspond to the strength with which each respective eigenmode participates in FC. Hence we define for each eigenmode *k* its “Graph Fourier Weight” (GFW) as *GFW*_*k*_ = |***Q***_*k,k*_|. Although this calculation requires the empirical FC matrix, it is not necessary to to use individual FCs, which would be impractical and circular. Instead, we found that the weights are reliably stable between individuals, hence they were pre-computed using average FC and SC matrices.

#### 3.3.3 Joint fitting algorithm

The model parameters *τ, α* are optimized per subject to maximize two objectives simultaneously: the spectral correlation and the FC correlation, as defined below. In this manner, we can obtain a unified model that reproduces both the regional power spectra and the second-order covariance statistics given by the frequency-resolved FC matrix (i.e. cross-spectral density). The cost function we used is therefore defined as:

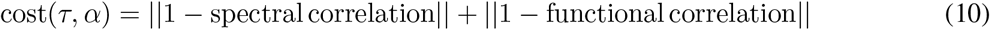

Functional correlation was defined as the Pearson’s correlation between the theoretical and empirical FC (only the upper-triangular portion thereof), while spectral correlation is defined analogously on the PSD. We implemented a constrained cost function minimization, available as the routine `fmincon()` in MATLAB version R2021b. The parameters *τ, α* were given lower limits 0 (to ensure positive values). We used default options for `fmincon()`, with 50 iterations and tolerance of 10^−6^ as stopping criterion.

### 3.4 Comparison to Randomized Structural and Functional Connectivity

To compare our proposed model to random structural networks, we constructed a null distribution by randomly sampling our dataset 1000 times and generating a randomized SC network using the Brain Connectivity Toolbox function `randmio_und_connected()` [62]. This function preserves the network’s degree distribution while ensuring the graph remains fully connected, which is highly significant for the Laplacian eigenmodes. To compare our proposed model to random functional networks, we again constructed a 1000 sampled null distribution by randomly permuting the BOLD time series region ordering, hence randomizing the FC matrix. We use the randomized SC to predict empirical FC, and we use the empirical SC to predict the randomized FC and spectra ordering. We compare all the resulting distributions by two-tailed t-tests.

### 3.5 Comparison to Benchmark methods

There is no current model that fits to and predicts the spectral content of fMRI. However, numerous models are available that predict FC, i.e. the covariance structure of the signal at zero lag. We compared the FC portion of our results against archetypal model-based approaches for the prediction of FC that explicitly involve the structural connectome (SC). Therefore we excluded spectral DCM method since they do not involve SC, but included both linear and non-linear SC-based graph models. We also excluded recent matrix algebraic methods like [31], on the grounds that it uses, and needs to infer, a large number of model parameters, e.g. a full *N* × *N* rotation matrix [31, 35]. Such models do not concord with the attribute of parsimony that is essential to our approach.

#### 3.5.1 Linear Algebraic Graph Model Comparison

Due to their emerging popularity and conceptual similarity, algebraic graph models (i.e. those that involve a transformation of SC eigenvalues) are highly pertinent benchmarks for our current proposal. Although many such methods are now available [35], we chose the two models which are both the earliest and the most parsimonious: the Network Diffusion Model [27] and Eigen-Decomposition Model [29]. Both models have similar parsimony to ours: 1 parameter and 3 parameters, respectively.

#### 3.5.2 Neural Mass Model Comparison

To compare proposed the SGM model to the more commonly reported generative simulations involving nonlinear neural dynamics, we implement the NMM model described by Breakspear and colleagues [63, 64] using the same model parameters previously used in [27] and [29]. A free coupling parameter(*c*) modulates the inter-regional coupling, and we evaluate the model over a range of *c* ({0.05, 0.1, 0.15, 0.2, 0.25, 0.3, 0.35}) and compare this model to SGM for fMRI using the optimal *c*.

## 4 Results

Our results are organized as follows. First, we establish that there is indeed interesting frequency content in rs-fMRI data, even in conventional recordings obtained at 2s TR, confirming prior work on coherence FC [14, 15]. We then demonstrate the power of the proposed model-based analysis of frequency-resolved fMRI by fitting the proposed SGM to empirical data. Our hypothesis is that even a global, spatially-invariant model with only 2 biophysical parameters should be able to predict conventional zero-lag FC and power spectra. We show this first on slower, conventional acquisitions obtained at our institution, since the vast majority of fMRI data are acquired this way. Of course, the true power of the approach would be better leveraged when fitting to fast acquisitions with a wider frequency range - this is our final result, and it is demonstrated on a high-quality, public fMRI dataset consisting of 50 healthy subjects.

### 4.1 FC has important spectral information

We first show that fMRI signal has frequency-dependent information content worth modeling. Figure 2A shows a set of cross-correlation signals *R*_*ij*_(*τ*) over a range of *τ*. The cross-correlation function is symmetric around 0, hence only the positive lags are shown. It is clear that various region-pairs give maximal correlations at non-zero lags, with many peaking in the range *τ* : 10 − 12*s*. Interestingly, region pairs whose zero-lag correlations are small or negative tend to produce high correlations at non-zero lags. Figure 2A also contains a histogram of the lags at which maximum correlation was achieved. The range of lags is typically within 12-15 seconds, and is widely distributed in that range. Figure 2B shows the 86 × 86 × *N*_*freqs*_ frequency resolved cross-spectral density (CSD) array: this corresponds to the spectral power of the cross-correlation function. Due to its evident frequency-selective distribution, it contains information beyond the conventional zero-lag FC, which corresponds to the integral over all frequencies of the CSD array. Since the full CSD volume contains large components wiht little signal, we carefully chose the most information-rich features from the full CSD: the power spectral density (PSD, defined as the diagonal of the CSD volume); and CSD at a specific frequency *ω*_0_ at which the pairwise sum of CSD has the highest value for a given subject. These features are illustrated in the figure.

### 4.2 Predicting FC and BOLD Spectra from SC using the Spectral Graph Model

Using SGM induced by the structural Laplacian matrix, we find that the model performs well, fitting simultaneously to zero-lag FC and BOLD fMRI spectra with mean (std) Pearson’s *R* = 0.59 (0.08) for FC and Pearson’s *R* = 0.70 (0.08) for BOLD spectra (Figure 3A). We additionally report optimal parameter values mean (std) for *α* = 0.80 (0.09) and *τ* = 1.96 seconds (0.56 seconds) (Figure 3B). In Figure 3C, we show three subjects’ example SC, empirical FC, and SGM-predicted FC, along with their individual Pearson’s *R* similarity; in Figure 3D, we show the same three subjects’ empirical and predicted BOLD spectra also with its associated Pearson’s *R* similarity. From these results, SGM clearly reproduces salient elements of both empirical FC and BOLD spectra, using a single model parametrization.

**Figure 3:**
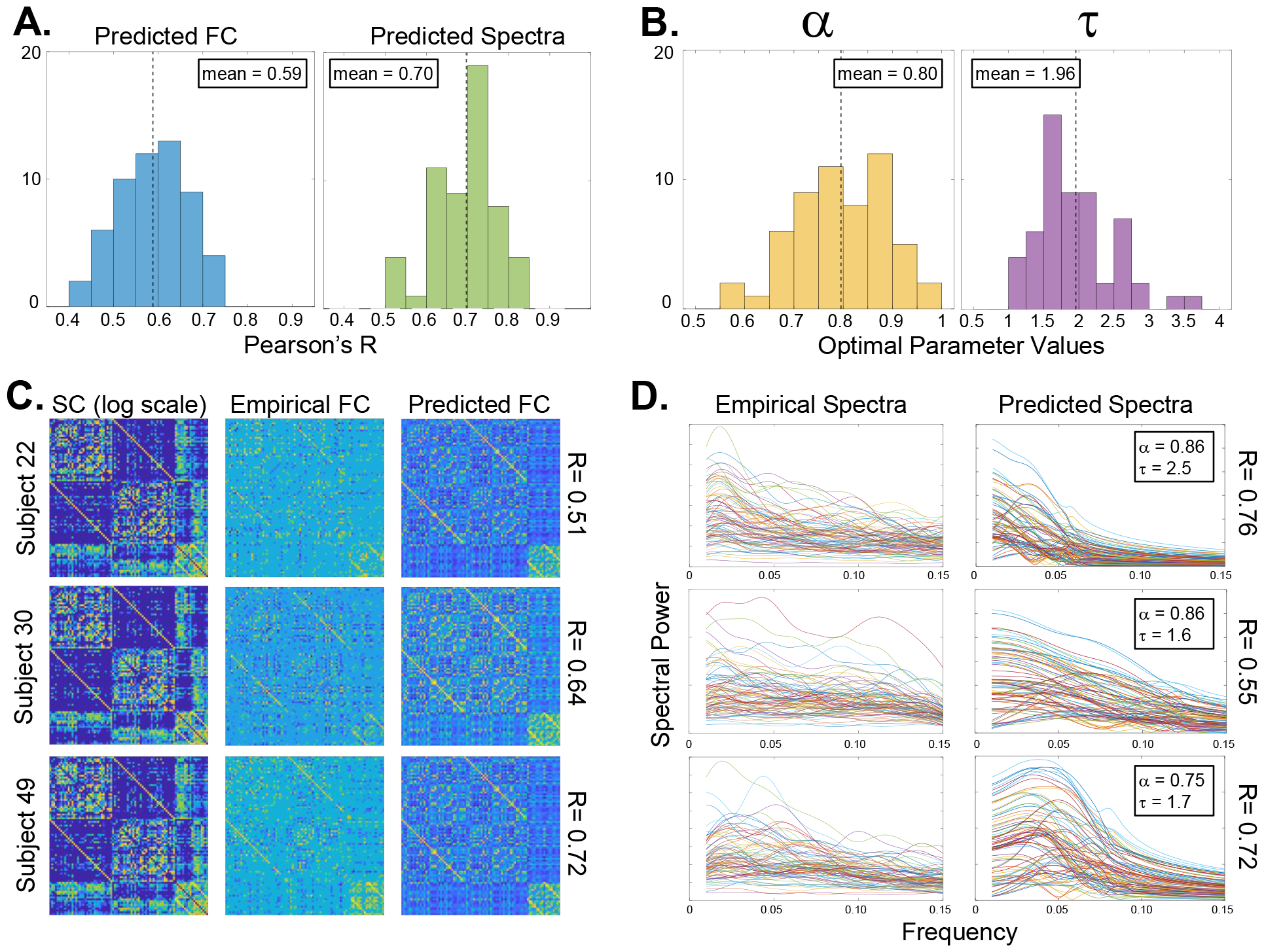
Results of SGM model fitting to UCSF fMRI dataset (*N* = 56). Histograms in panel **A** show the goodness-of-fit (Pearson’s *R*) between predicted and thresholded empirical FC at *ω*_0_, taking only the upper-triangular portion (see Figure 2**D**). The mean (standard deviation) *R* across all subjects was 0.59 (0.08). The second histogram gives the distribution of Pearson’s *R* computed between predicted and empirical PSD, averaged over all regions. The mean (std) *R* across all subjects for spectra was 0.70 (0.08). The distributions of optimally fitted parameters *α* and *τ* is shown in panel **B**. The mean (std) for *α* and *τ* Were 0.80 (0.09) and 1.96 seconds (0.56 seconds), respectively. Both parameters appear to fall within consistent, biologically-plausible ranges. Panel **C** contains empirical and predicted FC matrices from three representative subjects, along with their SC (log-scale for visualization). Panel **D** contains the same subjects’ PSD, both empirical and SGM-predicted. Since our goodness of fit is an average over all regions, the model spectra do not attempt to match outlier regions in the empirical spectra. These examples showcase the ability of a single SGM to simultaneously reproduce both FC and spectral features of individual subjects.

### 4.3 Demonstrating the power of SGM on wideband fMRI spectra from fast fMRI acquisitions

The vast majority of fMRI scans, like the previous results, have *TR ≥* 2 sec, hence in order to truly capitalize on frequency content we next applied the proposed model-based analysis to a public dataset of fast acquisitions (*TR ≈* 0.6 sec). As shown in Supplemental Figure S1, SGM for fMRI performed slightly better on this new frequency-rich data (mean(std) FC-R = 0.57(0.08); Spectral-R = 0.87(0.04) than our previous result. This not only supports the model’s capacity to account for higher frequencies, but also serves as independent confirmatory evidence analogous to the main results in Figure 3.

### 4.4 Effect of implementation choices

Several algorithmic and practical considerations affected the final technique demonstrated above. In this section we briefly explore the effect of these choices and provide empirical support for why they were chosen.

#### 4.4.1 Eigenmode Weighting using Graph Fourier Weights (GFW)

We weighted the model’s eigenvalues by GFW, which quantifies how much the eigenmode is expressed in the average FC’s projection matrix ***Q*** (Figure 4A). The group-averaged GFWs across the eigenspectrum are shown in Figure 4C. This reweighting of eigenvalues alters the eigenmode’s theoretical frequency response (Figure 4B), and substantially improves the model’s overall performance (Figure 4D). To our knowledge, the exact manner in which we have implemented eigenmode weighting has not appeared previously.

**Figure 4:**
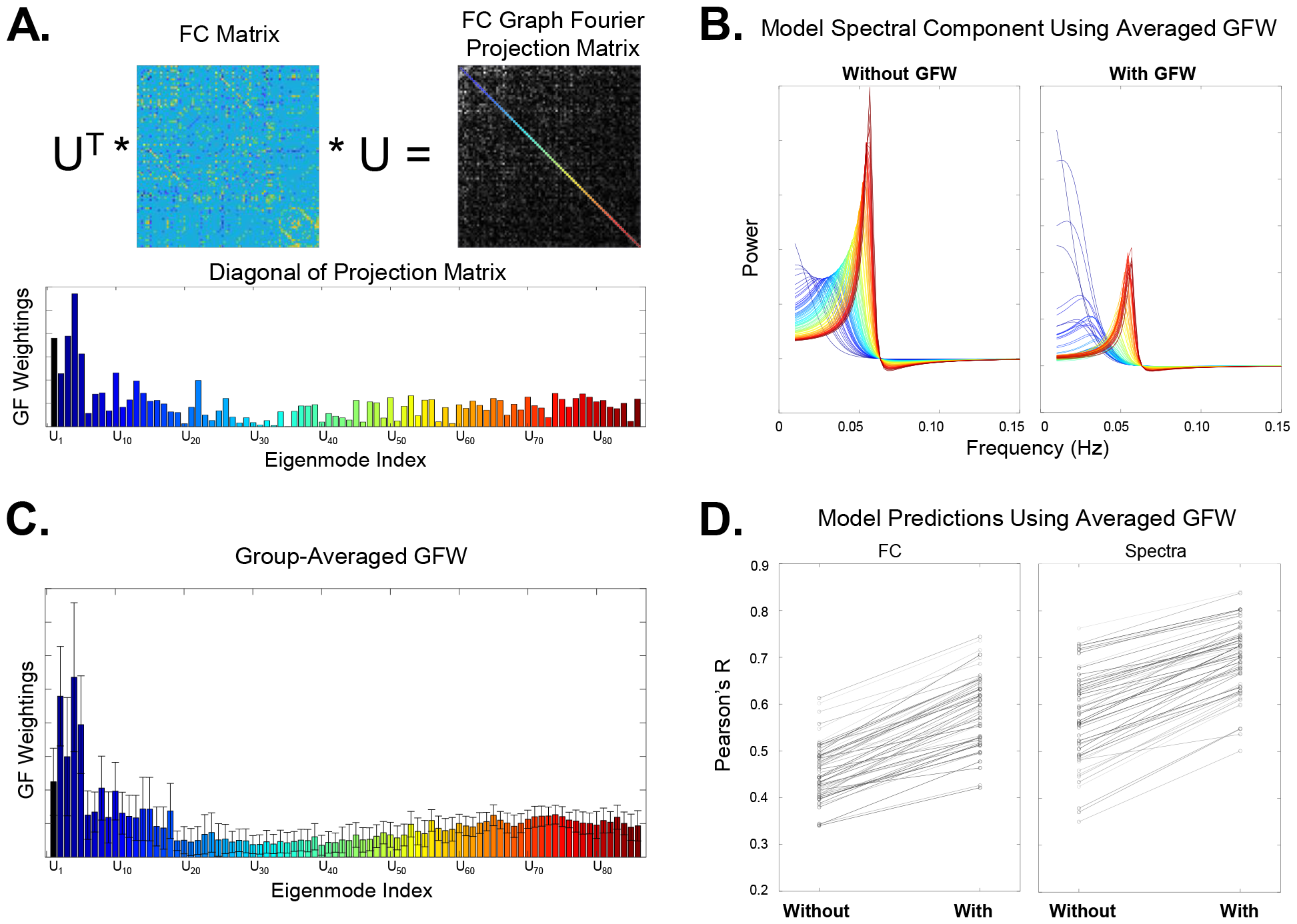
Eigenmode Weighting. The Graph Fourier projection matrix ***Q*** = ***U***^***H***^ *FC* ***U*** of a single subject is shown in panel **A**, clearly depicting the diagonal-dominance predicted by theory; here the diagonal is color coded by eigen-index. The diagonal entries’ magnitude, which defines our GFW weights, is shown in the accompanying bar chart. Panel **B** shows the impact of this reweighting on each eigenmode’s theoretical frequency response (inner terms in Eq 7). Panel **C** depicts the GFW obtained from all subjects; which are clearly similar to the individual subject’s GFW in **A**. Error-bars show one standard deviation above and below the mean. Henceforth the cohort-wide average GFW is used in all analyses. Panel **D** shows that the inclusion of GFW in the model of Eq 7 results in substantially and universally enhanced overall performance, given here by our chosen goodness of fit (Pearson’s *R*) between model and empirical FC and power spectra.

Of course, there are many alternative decisions we could have made regarding individual versus group-averaged eigenmode weights, as well as how to threshold the empirical FCs using the percolation approach. For completeness, Table 1 summarizes results for these different conditions. All results contained elsewhere in this paper comprised of the best condition set from this table, highlighted in **bold**.

**Table 1:** Evaluation of practical choices regarding thresholding and eigenmode weighting. We show SGM for fMRI results for various different choices that could have been made to evaluate the model. The three most fundamental choices were with respect to the percolation threshold, the Graph Fourier Weighing (GFW), and to adding the Interhemispheric and Regional Adjacency weights. For the percolation threshold and GFW, there were three options, to use subject-specific information, to use the group mean values, or to omit the modification. Because adding interhemispheric weights and regional adjacency information was uniform across all subjects, here we simply list ‘yes’ or ‘no.’ We have shown our final results above in **bold**. Unsurprisingly, the model performs much better when evaluating only the strongest (and therefore less noisy) FC matrix, as well as weighting the model’s eigenvalues to reflect the FC’s graph fourier projection. While the best results occur with all individualized information, for the reasons described above, we discuss results for group-averaged GFW.

| Threshold | GWF | Interhemi+Adj | FC R-val | Spectra R-val | $\alpha$ | $\tau$ |
| --- | --- | --- | --- | --- | --- | --- |
| Individual | Individual | Yes | 0.63 (0.07) | 0.70 (0.08) | 0.82 (0.08) | 1.94 (0.46) |
| **Individual** | **Group Mean** | **Yes** | **0.59 (0.08)** | **0.70 (0.08)** | **0.80 (0.09)** | **1.96 (0.56)** |
| Group Mean | Individual | Yes | 0.57 (0.06) | 0.67 (0.08) | 0.89 (0.07) | 1.81 (0.42) |
| Group Mean | Group Mean | Yes | 0.57 (0.06) | 0.67 (0.08) | 0.89 (0.07) | 1.81 (0.42) |
| None | Individual | Yes | 0.41 (0.05) | 0.69 (0.08) | 0.81 (0.08) | 2.15 (0.50) |
| Individual | None | Yes | 0.46 (0.06) | 0.58 (0.10) | 0.84 (0.14) | 2.02 (0.43) |
| None | None | Yes | 0.34 (0.04) | 0.59 (0.10) | 0.78 (0.16) | 2.19 (0.46) |
| Individual | Individual | No | 0.50 (0.09) | 0.64 (0.08) | 0.74 (0.17) | 2.51 (0.67) |
| None | None | No | 0.25 (0.04) | 0.55 (0.11) | 0.21 (0.15) | 2.97 (0.62) |

### 4.4.2 Alternate definitions of PSD and FC

The covariance and Fourier formalism given in the Theory section accommodate multiple definitions of both what constitutes FC, and what constitutes PSD, depending on the underlying assumptions. We have thoroughly evaluated the effect of these choices, summarized in Table 2. In these explorations we used the best condition from Table 1, using individual thresholds for FC and group-mean eigenmode weighting.

**Table 2:**
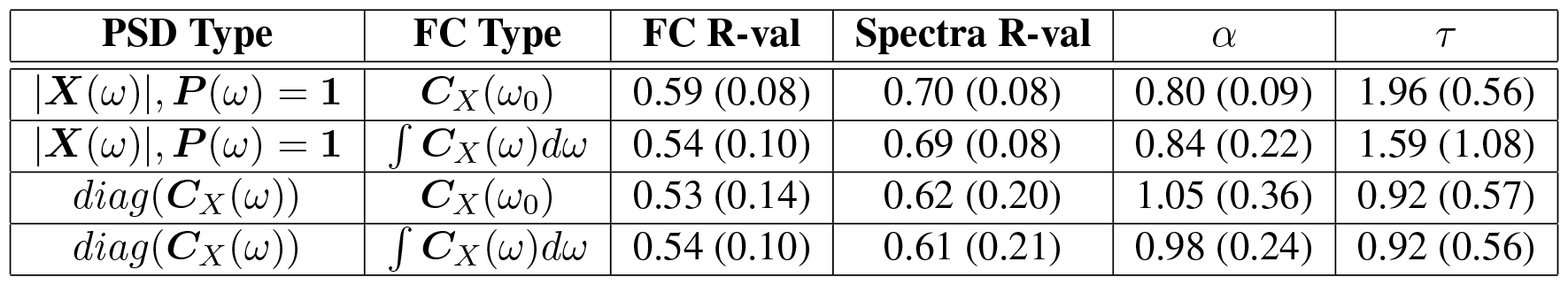
Evaluation of alternate definitions of PSD and FC. We show SGM model fits under different definitions of Power Spectral Density (PSD), and of FC. FC was defined either at the maximum-amplitude frequency (*ω*), reflecting the frequency with the most signal energy; or as an average across all frequencies, reflecting zero-lag FC. The SGM model performs well under all these condition, but the direct PSD under uniform input power, and FC at the highest-amplitude frequency, give the best predictions (top row).

The main results of this paper in Figures 3 and S1 employed the maximum-signal slice of the CSD volume for FC, i.e. ***C***_*X*_(*ω*_0_). However, the formal definition of zero-lag FC, given by ∫***C***_*X*_(*ω*)*dω*, also gives a very good match to empirical data, with mean (std) Pearson’s *R* = 0.54(0.10) for FC and Pearson’s *R* = 0.69(0.08) for BOLD spectra. This is shown in the first 2 rows of Table 2. We did not have an *a priori* preference of zero-lag FC over specific FC evaluated at *ω*_0_ - but from a practical point of view the latter was found to be more reliable to fit and gave slightly better results. In similar vein, both the formal PSD, as the diagonal of the CSD, i.e. *diag*(***C***_*X*_(*ω*)), and the direct signal power under an explicit input, |***X***(*ω*)|, when ***P*** (*ω*) = **1**, produced good results. Table 2 shows that the direct PSD |***X***(*ω*)| in fact does slightly but consistently better than *diag*(***C***_*X*_(*ω*)) at fitting to empirical BOLD power spectra.

#### 4.4.3 Comparison with 3-parameter SGM

As noted in the Model section, our model specification requires only 2 parameters: *α* and *τ*, the latter being shared by both the Gamma-shaped cortical response function as well as by the network spread process. Our motivation was essentially parsimony; yet there was no special reason why the two processes could not be characterized by different time constants. To empirically support our choice we performed an empirical comparison between the two versions. We split the singular time constant *τ* into two: *τ*_1_ and *τ*_2_, the latter denoting the cortical response time. The results are shown in Supplementary Figure S2 and compared against the original 2-parameter model. For these comparisons we initialized both time parameters to the same value and gave them an identical range. As it transpired, the two constants meaningfully diverged from the singular version, yet the end model was no better than the original, more parsimonious one. We deemed the net benefit of adding the new parameter to be negative; henceforth all results are presented for the 2-parameter model.

### 4.5 Benchmark Comparisons with Previous Models

We sought to assess how SGM for fMRI performed in comparison to previous linear SC-FC models implemented on the same dataset. Since existing models can only predict FC and not the PSD of fMRI signal, we used only the FC component of our predictions in these comparisons. We observed that SGM obtains better fits to individual FC compared to both previous linear graph models, the Network Diffusion Model [27] and the Eigen-Decomposition model [29] (Figure 5). Then we assessed the performance of the connectome-coupled Neural Mass Model [63, 64], whose Pearson’s *R* was centered around 0.44 – lower than above linear models and far lower than the proposed SGM model. Using a one-way analysis of variance (ANOVA), we found that there was a significant group-wise difference (*p <<* 0.01) between the SGM and all other benchmark models. Importantly, SGM showed better performance compared to all alternative models.

**Figure 5:**
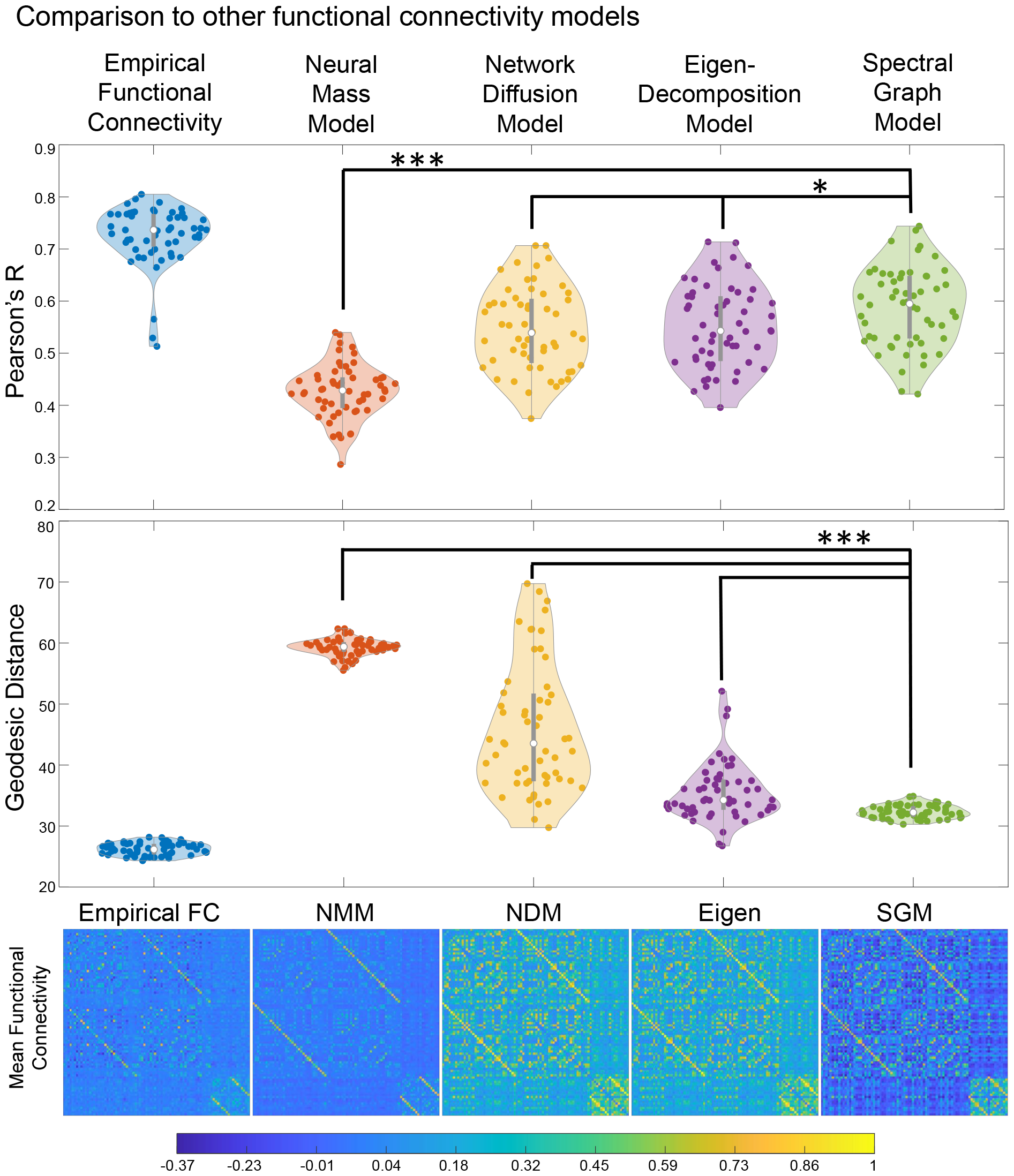
Benchmark Model Comparisons. **Top row**: In a violin plot, we compare SGM for fMRI (green) to alternative FC prediction models, namely a Neural Mass Model (red) [63, 64], the Network Diffusion Model (yellow) [27], and the Eigen-Decomposition model (purple) [29], using Pearson’s R as goodness of fit (∗ : *p* < 0.05; ∗ ∗ ∗ : *p* << 0.001; 1-way ANOVA: *p* = 7 ∗ 10^−26^. To provide additional context to these numbers we also show the distribution of the inter-subject similarity in empirical FC, using cohort-wide mean FC as reference. **Middle row**: We report the geodesic distance as an alternate goodness of fit, considered more appropriate for semipositive definite matrices. Under the geodesic measure the proposed SGM model performs far more significantly^2^b^2^etter compared to benchmark methods than under the conventional correlation measure. **Bottom row**: We show the mean predicted FC across all subjects for each model (1-way ANOVA: *p* = 2.1 ∗ 10^−63^).

### 4.6 Comparisons with Null Models Using Random Networks

To ensure that SGM for fMRI goodness of fits are specific to the structural network, we sought to test whether randomizing the SC matrix diminished SGM’s capacity to fit to empirical FC and BOLD spectra. Indeed, we found that similarity distributions using randomized samples of subject-specific SC were significantly different than the empirical SC fit to FC (*p* = 0) and to BOLD spectra (*p <<* 0.001) (Figure 5A-B).

We also sought to evaluate whether SGM for fMRI could predict FC derived from a time series obtained by randomly reordering regions (***T*** ^*rand*^) while maintaining the empirical SC. We found that randomly shuffling the ROI time series again destroyed SGM’s ability to predict FC (*p* = 0) while not impacting the prediction of BOLD spectra (*p* = 0.85) (Fig 5C-D).

#### Interpretation

The results from the null-models require careful assessment. The predicted FC from a randomized SC is clearly separated from the predictions using empirical SC, yet the spectral predictions for randomized versus empirical SC are significantly different but not as well separated. It is likely that SC randomization destroys the spatial patterns and region order inherent in FC, while the spectral content is less affected since in theory the random graph can encompass eigenmodes and eigenvectors that may be nearly sufficient to produce a desired set of spectra. This is observed even more clearly in the second randomization, that of time series regional order. Here again, the implied FC is destroyed due to region reordering, but the spectra are preserved (albeit with a random reassignment to regions). This implies again that the predicted model’s spectra do not possess strong regional specificity.

## 5 Discussion

### 5.1 Key findings

There is a need for innovation in new-generation analysis tools for rs-fMRI focused on overcoming current gaps. In particular, we identified a) the opportunity to use rich spectral content available in modern multi-band fast acquisition techniques; and b) the opportunity to relate fMRI analysis products to biophysically interpretable parameters, which might one day become useful as biomarkers of task, cognitive or disease states. The proposed spectral graph model uses the brain’s structural network to generate a synthetic fMRI signal; in this study it is implemented as a general purpose yet biophysically interpretable tool to analyze wideband fMRI and its induced FC simultaneously. It does so with the simplicity and parsimony of spectral graph theory, using only two global parameters that between them accommodate the variability inherent within healthy subjects’ FC and spectra: *α*, which accounts for global coupling, and *τ*, which accounts for signal propagation speed. The model naturally decomposes into a sum over the eigenmodes of the structural graph Laplacian. Remarkably, the model admits closed-form solutions of the regional signal’s frequency spectra as well as frequency-dependent FC - a useful departure from existing coupled neural systems that require lengthy numerical simulations. A key refinement of the approach involved *a priori* weighting of the Laplacian eigenmodes such that only those eigenmodes that contribute to the resting-state fMRI signal patterns in the entire cohort were retained in the summation (Figure 4).

We reported excellent capacity of the model to predict empirical power spectra as well as FC, on both conventional acquisitions (Figure 3) and fast multi-band acquisitions (Figure S1). We found that frequency-resolved SGM produced significantly improved predictions of FC over previous linear and non-linear SC-FC models, significantly exceeding them in both correlation and geodesic measures of performance (5). Most strikingly, SGM gave far better fits than a coupled system of non-linear neural masses, supporting the idea that macroscopic neurophysiological data on a graph can be sufficiently modeled with linear metrics, and nonlinear methods may not be required for problem of such scale [65, 66]. We demonstrated that SGM’s predictions are specific to the SC network topology by comparing to a null distribution of randomized SC matrices. Our results were far better than could be expected by chance (Figure 6). In the following sections, we discuss the implications of this spectral graph model and contextualize these results against the current literature.

**Figure 6:**
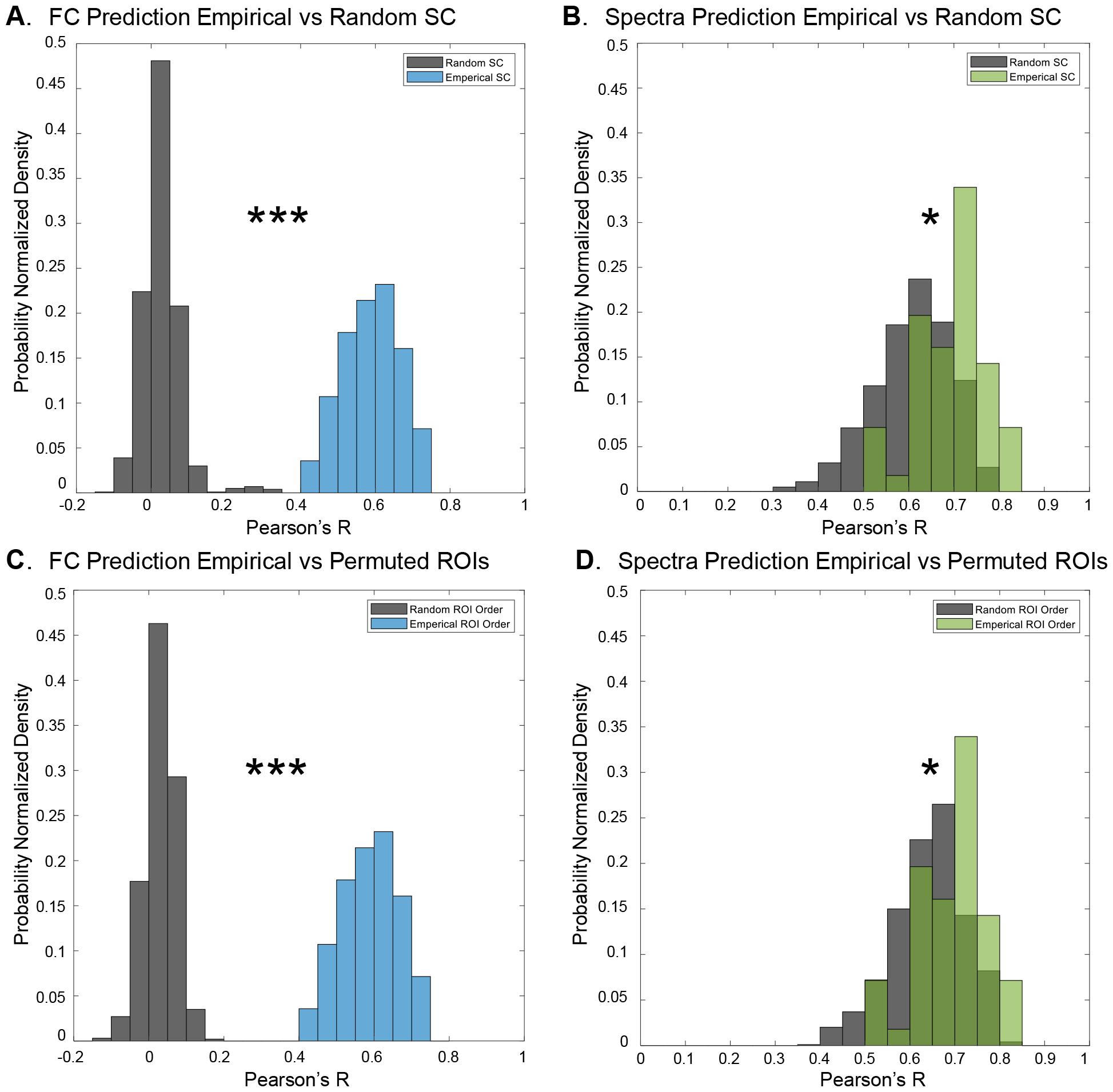
Validation with Null Models. We compare SGM for fMRI to randomized SC and time series. We predict functional connectivity (A) and BOLD spectra (B) using a null-model randomized SC that preserves the connectomes’ degree distribution while ensuring it does not become disconnected (which would significantly change the Laplacian eigenmodes). In both cases, the randomized SC (shown dark gray) creates ‘optimal’ fits that are significantly worse than the empirical SC (colored) (\**p <* 0.001; \*\*\**p* = 0). We further predict FC (C) and PSD (D) after randomizing all functional ROI labels to generate a randomized FC matrix while fixing SC. Again, randomly permuting ROI time series labels results in significantly worse model performance, which is especially pronounced for the functional connectivity.

### 5.2 Frequency-resolved SGM can accommodate a rich repertoire of spatial-spectral patterns, tunable with two biophysical parameters

Our model was able to access most observable configurations of spatial and spectral patterns seen in real rs-fMRI data by tuning only two global and biophysically-meaningful parameters: coupling strength and neural time constant. That such a parsimonious and analytical model can produce a frequency-rich repertoire of activity may be a novel contribution to the field. As observed from Figures 3 and S1, parameter regimes typically recruit and “steer” multiple eigenmodes in such a manner that a small number of them can reproduce signal spectrum and FC at any relevant frequency. Possibly, this points to an essential characteristic of real brain activity, which is thought to accommodate a large repertoire of microstates and their concomitant spatial patterns. Available literature on wide-band frequency brain recordings demonstrates that different fMRI functional networks are preferentially encapsulated by different higher-frequency bands via phase- and amplitude-coupling [67, 68]. Phase and amplitude coupling of oscillatory processes in the brain is evidently important for the formation of coherent wide-band frequency profiles of brain recordings and processing of information [69–71, 67, 68]. Historically, such processes could be modeled via large scale non-linear simulations of oscillatory neural activity [64, 72, 73]. In contrast, our approach does not require high-dimensional model parameters nor extensive simulations. This raises the possibility that complex behavior may be achievable by simple and parsimonious mechanisms.

The present computational study is not intended to explore the neural mechanisms that might control the model’s biophysical parameters. Nevertheless, several possibilities may be considered. Coupling strength*α* is a direct scaling of white-matter excitatory long range connections between neural populations in the brain. Parameterization of coupling strength between distant brain regions via the connectome is ubiquitous in connectivity based models of BOLD fMRI [74, 64, 68, 27] and electroencephalography activity [75–77]. Furthermore, pathological FC patterns as a result of disconnections in the brain can be reproduced with decrease in coupling strength [78, 79]. The other tunable parameter in our model, neural time constant *τ*, is the aggregate lumped time constant that captures in a single number the overall effect of membrane capacitance, conductance speed, delays introduced by the dendritic arbors of the ensemble of neurons within a region, and the local processing delays of signals within the region. It also incorporates the convolution of the neuronal signal with the hemodynamic response pertaining to the BOLD signal. While the individual components of these processes are likely complex and challenging to infer in a model such as this, their overall lumping into a single time constant *τ* enables us to capture emergent behavior without breaking the essential parsimony of the generative model. We empirically investigated the possibility that an additional time constant would improve the model further, but this was not the case; see Figure S2. Hence, it may be deemed that the proposed model is sufficiently parsimonious.

### 5.3 Relationship to existing approaches

We review two types of prior models relevant to current work: graph models and generative models.

#### 5.3.1 Graph models

Recent graph models involving eigen spectra of the adjacency or Laplacian matrices of the structural connectome have greatly contributed to our understanding of how the brain’s structural wiring (SC) gives rise to its FC. Such models typically assume that SC and FC are not independent entities, yet their relationship is not direct [64]. In addition to connection strength between regions, metrics such as anatomical distances [80], shortest path lengths [81], diffusion properties [29, 82], and structural graph degree [83] were also found to contribute to the brain’s observed functional patterns. Eigenmodes and eigenvalues of a network’s adjacency or Laplacian matrix express an ordered, low-dimensional, representation of that network’s topology. Eigenmodes are thus conceived to represent *spatial* frequencies of the underlying network—the fundamental spatial patterns contained within the network’s topology [84, 29, 33]. Reassuringly, these SC eigenmodes are reproducible, with the first few lowest “spatial frequency” patterns being similar across different atlas parcellation resolutions, between different human subjects, and between two scans from the same subject [85]. The success of eigen-mapping methods has been attributed to SC and FC sharing fundamental features in their eigenspace [29, 86]. A recent extension introducing transmission delays and a “complex Laplacian” showed that different sets of eigenmodes contributed in varying degrees to canonical RSNs [87]. Higher-order walks on graphs, involving a series expansion of the graph matrices, have also been quite successful [32, 31]. The diffusion and series expansion methods are themselves closely related [88], and almost all harmonic-based approaches may be interpreted as special cases of each other, as demonstrated elegantly in recent studies [35, 34]. Yet another mapping between FC and SC eigenvalues via the Gamma function was demonstrated [56].

In the Theory section we were able to show mathematically that the eigen-mapping model central to these studies is essentially a special case of the current generative model, reducing to the former when the neural response function is removed and the input power is i.i.d. Gaussian. The key differentiator however is that prior graph models have not considered the spectral content of the signal, and in fact do not have the ability to give non-trivial spectra whatsoever. Our concept of *a priori* learning and weighting the importance of the eigenmodes that go into the SGM model (Figure 4) is also inspired from prior work. Indeed, taking projections of SC eigenmodes into a functional space is a simple, powerful means to identify eigenmode weightings. A recent study has used the signal’s eigenmode projection as a form of filtering [60].

#### 5.3.2 Generative models

While keeping the parsimony of eigen-mapping graph models, our approach adds an explicit signal model with frequency content accommodating oscillatory frequency and phase shifts between brain regions. However, the SGM signal generation model is not the only example of generative modeling in fMRI analysis. In fact, many such generative models exist, under two broad forms: a) dynamic causal modeling with an unknown effective connectivity matrix [16] to fit a multivariate autoregressive process to the time series [89, 90] or complex fMRI cross spectra (“spectral DCM”) [16, 18]; or b) neural mass model (NMM) simulations, which use coupled non-linear NMMs, typically simulated first to much higher (up to 40 Hz) frequencies, then down-sampled to BOLD frequency ranges via a Hilbert envelope [91, 26]. Each method is briefly described below and compared against the current approach.

##### DCM and VAR

While the goal of VAR and DCM models in fMRI analysis is similar to our proposed model that makes model inferences about FC, the two frameworks are vastly different in terms of approach and dimensionality. DCMs examine the second order covariances of brain activity [92, 93], and it is only recent works with spectral and regression DCM models that have expanded the model coverage to the whole-brain scale [94, 95, 18]. Since they are not designed to be biologically grounded in the underlying connectome, DCM models have many more degrees of freedom compared to our work because they parameterize for different interactions within and between brain regions.

##### Neural mass models

An effective way to address higher frequency-information is to employ structural connectivity to couple anatomically connected neuronal assemblies [96, 97].Numerical simulations of such neural mass models (NMMs) provides an approximation of the brain’s local and global activities, and are able to achieve moderate correlation between simulated and empirical FC [98, 72, 99, 64, 76]. These coupled NMMs are perfectly capable of producing high-frequency content, but their existing use in fMRI modeling tends to focus on reproducing zero-lag FC exclusively and not on generating the fMRI signal PSD at non-zero frequencies. Moreover, these models rely on lengthy and expensive time series simulations of local neural masses to derive dynamical behavior, which are then used to generate effective connectivity matrices. Such approximations through stochastic simulations are unable to provide a closed form solution; their inference requires a large number of model parameters and exacerbates and inherits interpretational challenges. It is noteworthy that SGM for fMRI may be considered a highly-simplified and linearized NMM, but with a focus on the coupling between remote neural masses rather than detailed modeling of those masses themselves. Despite only fitting to FC (and not spectra as SGM does), the coupled NMM we implemented here was unable to produce results on FC comparable to ours, as demonstrated in Figure 5E. By avoiding large-scale simulations of neuronal activity, our approach overcomes these practical challenges and aids biological interpretability.

The modeling of higher frequencies is more common in the context of MEG data. A signal generation equation similar to ours was previously proposed for capturing higher frequencies obtained from MEG [100]. Their context was different (high MEG frequency regime) and their model involved local excitatory and inhibitory neuronal subpopulations and phase lags arising from axonal conductance - elements which are not pertinent to the fMRI regime. Despite several studies of associations between fMRI and MEG FC and phase amplitude coupling, there is not yet a single convincing model that can simultaneously capture both regimes.

### 5.4 Clinical applications leveraging ease of inference

Any analysis tool must eventually find purpose in practical and clinical domains. Since the proposed technique is highly parsimonious and admits closed-form solutions, it presents an unrivalled opportunity: easy and straight-forward fitting to real data, over any and all frequencies of interest. As shown in Figures 3, SS1 and 5, the model achieves excellent fits to individual subjects, quantitatively superior to FC predicted by prior linear models, and additionally fits to regional power spectra which no current model can achieve. Our work can therefore find direct applicability in many clinical and neuroscientific contexts where predicting functional patterns from structure is important [101, 22], particularly in cases of epilepsy [102, 30], stroke [82, 103], Schizophrenia [56, 13] and neurodegeneration [104].

### 5.5 Limitations of this study

Several assumptions and limitations are noteworthy. Our theorized model relies on a uniform neural time constant throughout the brain. In reality, the amount of myelination and synaptic strength varies greatly in the brain, as do the number of neurons and their local electrophysiological properties. However, this strong assumption did not adversely affect the model’s predictive ability, which is similar or higher than currently reported in the literature. Certainly, predictive power could be improved by the use of regionally varying model parameters, but the current approach has the inestimable benefit of a low dimensional and interpretable model. A linear model like SGM is limited in the repertoire of dynamics that it can produce. Interestingly, a recent comparison [105] demonstrated that linear models predicted resting fMRI time series better than non-linear models. They argued that this may be because of the linearizing affects of macroscopic neuroimaging due to spatial and temporal averaging, observation noise, and high dimensionality, indicating that a linear model may be sufficient to capture the observed fMRI dynamics.

The functional data used in this study had a 2-second TR temporal resolution, which limits the frequencies resolvable to the Nyquist limit of 0.25 Hz. Yet, most fMRI data are collected with 2-second TR, and conventional resting-state analysis doesn’t explore frequencies above 0.1 Hz, hence presented temporal resolution is appropriate given our aims. We used a gradient-descent optimization method, which is most effective in convex optimization. We expect that future implementations with non-convex Bayesian inference that generate parameter estimate posteriors can provide improvement. Further, our cost function is based on Pearson’s correlations between predicted and empirical FC and spectra, which causes our fits to reproduce broad similarities in spectral and FC shape rather than absolute scaling. Next, while weighting eigenmodes constitutes a clear improvement and novelty, this requires prior access to a group-level SC-FC data. Therefore our results could be strengthened in future studies by applying population averages from a separate large cohort.

## 6 Acknowledgement

This work was supported by NIH grants R01EB022717, R01AG062196.

## 7 Code & Data Availability

Code for SGM for fMRI has been made available at https://github.com/benjaminsipes/SGMforfMRI along with group-averaged SC. The full dataset can be made available upon request.

## 8 Supplementary Material

**Figure S1:**
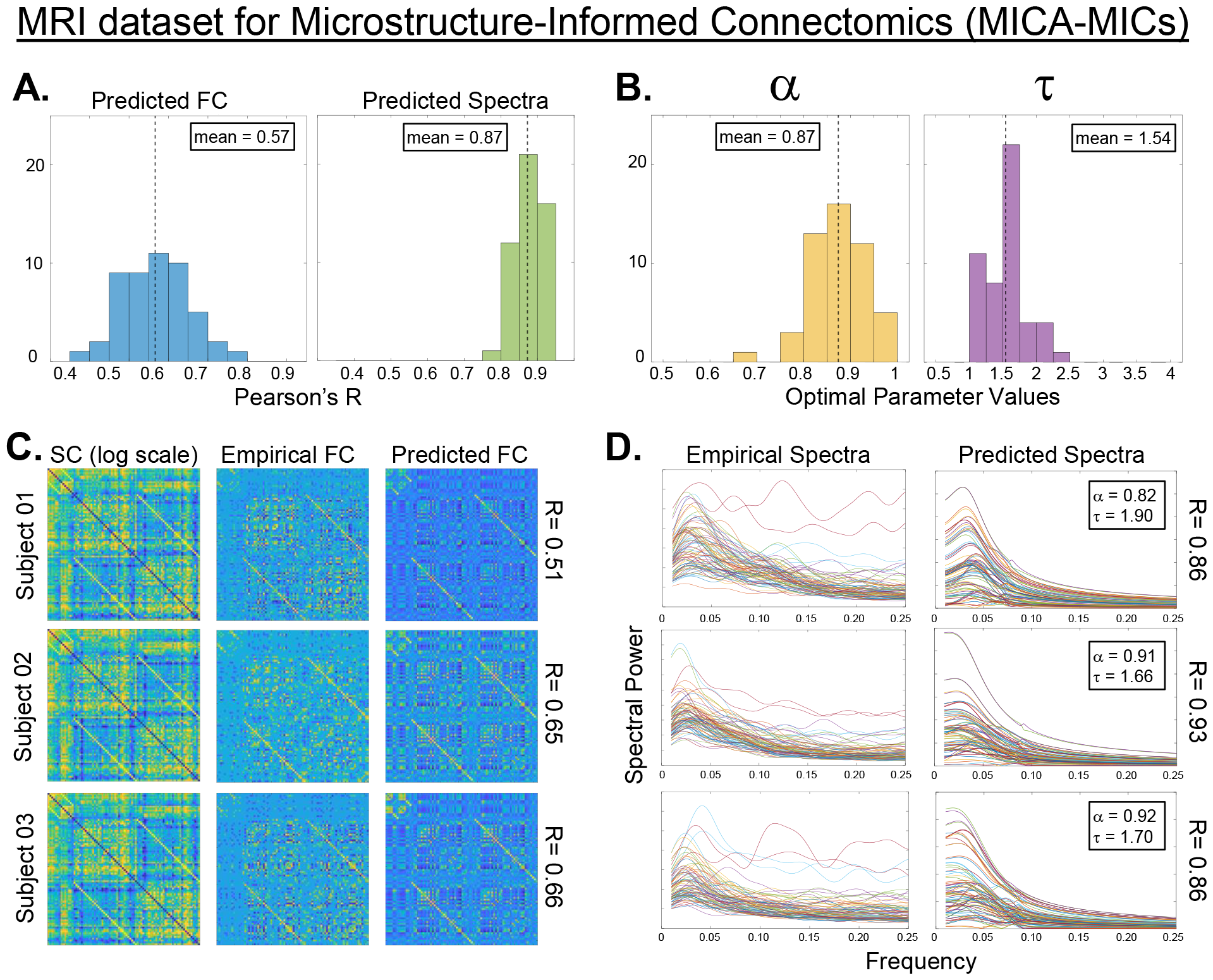
Replication with the high bandwidth public MICA fMRI dataset. (*N* = 50), analogous to Figure 3. The SGM model performed equally well in reproducing the faster and higher bandwidth MICA data (*TR* = 0.6 sec). Compared to the main UCSF study results, the histograms in panel A of goodness of fit (Pearson’s *R*) are similar, although the fitted model parameters in panel B are more narrowly-distributed.

**Figure S2:**
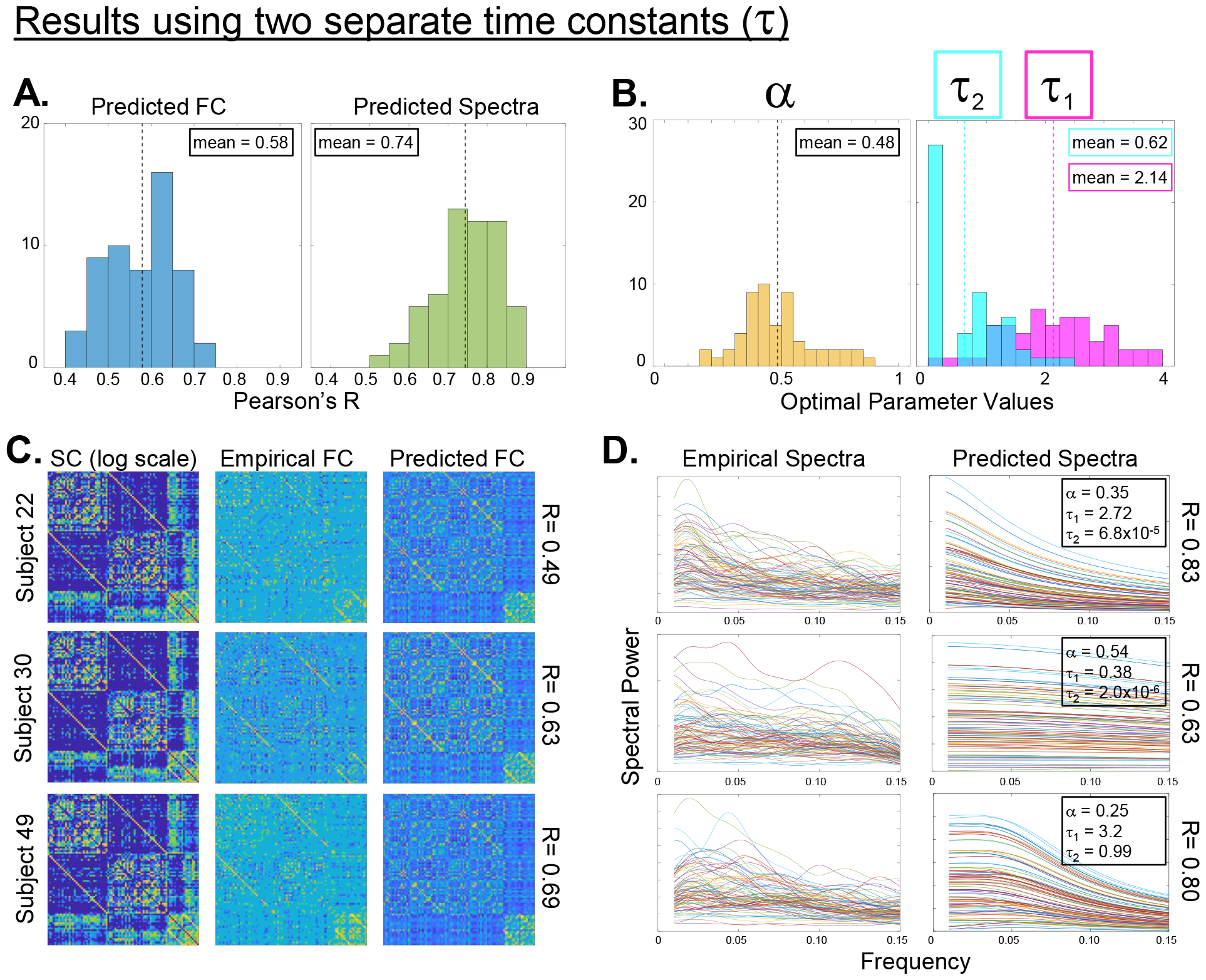
Model fitting on UCSF data under the 3-parameter SGM. We split the singular constant *τ* of the main model into two: *τ*_1_, the network diffusion time and *τ*_2_, the cortical response time. Both *τ*_1_ and *τ*_2_ received identical range and initialization. The results of this new model are presented here closely following the main Figure 3. While the fitted values of *τ*_1_ and *τ*_2_ meaningfully diverged from each other and from the singular version *τ*, yet the end model was no better than the original, more parsimonious one.

